# Versatile electroporation protocols enable reproducible CRISPR-RNP delivery across multiple primary mouse cells of the hematopoietic lineage

**DOI:** 10.64898/2026.01.27.702081

**Authors:** Ju Hee Oh, Lucy F. Yang, Erin Tanaka, Lauar de Brito Monteiro, Dasol Wi, Anne-Sophie Archambault, Ramon I. Klein Geltink

## Abstract

Genetic engineering of primary hematopoietic cells is essential for mechanistic immunology studies, however the development of cell-based therapies yet remains constrained by two major factors: the fragility of many primary lineages and challenges of viral delivery platforms that are costly, time-intensive, and biologically confounding. Here, we optimized scalable, non-viral CRISPR–Cas9 electroporation workflows using the ExPERT platform across three primary mouse hematopoietic cell types: OT-I CD8⁺ T cells, bone marrow–derived macrophages (BMDMs), and hematopoietic stem cells (HSCs). In activated OT-I CD8⁺ T cells, two electroporation programs supported high-efficiency mRNA and RNP delivery with minimal impact on cell viability or proliferative capacity, with subtle activation-state–dependent sensitivity at higher energy settings. Extending optimization to myeloid and stem cell lineages, we found that BMDMs maintained high viability following electroporation, and a high-performing electroporation program supported robust RNP delivery and efficient target gene knockout while preserving macrophage differentiation. In HSCs, the same program enabled consistent RNP delivery, sustained viability, and reproducible gene knockout. Together, these findings establish ExPERT electroporation as a robust, reproducible, and modular platform for non-viral genome editing across primary mouse hematopoietic lineages, lowering barriers to rapid genetic perturbation for both discovery and translational applications.

## Introduction

Primary hematopoietic cells are central to both mechanistic discovery and the development of cell-based therapies, yet genetic manipulation of these cells remains constrained by limited cell viability post-editing and reliance on resource-intensive viral delivery methods^1,2^. Establishing scalable, non-viral genome editing strategies that preserve cellular function is therefore essential for advancing both basic and translational research. Key hematopoietic populations of interest include CD8⁺ T cells, macrophages, and hematopoietic stem cells (HSCs), which collectively underpin disease modeling, therapeutic engineering, and immune function,

CD8^+^ T cells play a central role in adaptive immunity by contributing to host defense and anti-tumour responses^3,4^. Harnessing these CD8^+^ T cells has become a cornerstone of modern cancer immunotherapy. Among the most promising strategies is adoptive cell therapy (ACT), which involves the isolation, *ex vivo* expansion and/or engineering, and reinfusion of autologous T cells with enhanced specificity and effector capacity. ACT, including chimeric antigen receptor (CAR) T cell therapies, harnesses genetically modified T cells to recognize and eliminate tumour cells^5–7^. The first two CAR-T products approved by FDA were produced by gamma-retroviral (Yescarta)^8^ or lentiviral (Kymria)^9^ gene transduction where synthetic receptor constructs are packaged using double-stranded viral elements in mammalian cells, followed by concentration of the viral particles which can be used to effectively deliver the CAR encoding DNA into T cells^10^. Similarly, viral constructs can be used to encode specific short hairpin RNA (shRNA) molecules to induce RNA interference, leading to mRNA degradation and translational repression^11^. However, application of viral gene or shRNA transduction is limited by high manufacturing costs, complex production pipelines, and off-target editing safety concerns, including semi-random genomic integration and insertional mutagenesis^12,13^. These challenges present opportunities for optimization and improved scalability in both basic biological studies and the application of findings to improve immune cell therapies.

While T cells have are central to cancer immunotherapy, macrophages are increasingly recognized as promising effector cells. Playing an important role in innate immune activation and phagocytic clearance^14,15^, macrophages also possess protective and pathogenic functions in bone marrow transplantation^16^. Bone marrow-derived macrophages (BMDMs) from mice are widely used to study macrophage biology owing to well-established differentiation protocols that yield large cell numbers^17^. However, significant technical challenges create difficulty in achieving effective gene disruption in BMDMs and hamper their applications. This difficulty stems from the heightened sensitivity to exogenous nucleic acids, including viral vectors, which can induce innate immune activation^18,19^.

Beyond differentiated myeloid cells like BMDMs, hematopoietic stem cells (HSCs) represent another key hematopoietic population in which precise genetic manipulation is both highly desirable and technically challenging^20,21^. HSCs serve as a valuable source for the generation of various myeloid lineage immune cells and as such broadly used in basic research. Clinically, HSCs are central to many therapeutic strategies including autologous transplantation of genetically corrected cells for inherited disorders and allogeneic transplantation to reconstitute the immune system in patients with treatment-resistant malignancies^22^.

Genetic engineering of primary cells has historically relied on viral overexpression, shRNA-mediated knockdown or nuclease-based platforms such as transcription activator-like effector nucleases (TALENs). These approaches are limited by scalability, genomic integration, and cell-type-specific toxicity^23–25^. The CRISPR (<u>c</u>lustered <u>r</u>egularly interspaced <u>s</u>hort <u>p</u>alindromic <u>r</u>epeats)–Cas (CRISPR-associated) system has vastly improved the possibilities of targeted genetic engineering strategies^26,27^. This advance was enabled by the identification of its core functional components: the Cas9 nuclease^28^, a CRISPR RNA (crRNA) that specifies target DNA recognition^29^, and a trans-activating CRISPR RNA (tracrRNA) which can be combined with crRNA into a single guide RNA (sgRNA) to simplify delivery and editing workflow^30,31^.

Compared with shRNA-based knockdown, CRISPR-mediated gene deletion has emerged as a powerful and precise approach for dissecting function of genes and the encoded proteins^32,33^. The recent clinical approval of Casgevy, which uses non-viral CRISPR–Cas9 editing of human HSCs by using MaxCyte ExPERT platform, demonstrates that scalable RNP delivery systems are clinically feasible ^34^. Consistent with this, electroporation-based gene editing in human macrophages has shown strong translational promise^35^. However, adopting human cell engineering protocols to primary mouse immune cells remains challenging, limiting cross-species mechanistic and preclinical studies. Moreover, despite the molecular precision of CRISPR, viral delivery in sensitive primary cells is often constrained by limited payload capacity and prolonged nuclease expression, which can increase the risk of off-target editing. Together, these limitations underscore the need for optimized, non-viral delivery strategies that preserve cellular viability and function.

To address these limitations, non-viral gene delivery methods such as electroporation (EP) have emerged as promising alternatives for introducing CRISPR-Cas9 ribonucleoproteins (RNPs) and mRNA into primary cells. EP introduces editing machinery transiently, avoiding genomic integration and unintended off-target effects, and is compatible with good manufacturing practice (GMP) standards, making it an attractive platform for clinical translation^36,37^. Despite these advantages, efficient EP in primary cells remains non-trivial.

Primary cells are sensitive to alterations in culture media components such as serum, cytokines, growth factors, and editing efficiency is strongly influenced by cellular activation/differentiation states^38^. Effective gene editing therefore requires careful optimization of EP timing and parameters to balance high editing efficiency and cell viability, and preservation of the cell’s original and the capacity for differentiation into effector T cells and macrophages.

In this study, we optimized MaxCyte ExPERT protocols for CRISPR-Cas9 RNP-mediated gene editing across multiple primary mouse cell types, OT-I transgenic CD8⁺ T cells, BMDMs and HSCs. In CD8^+^ T cells, we evaluated and compared two protocols, originally designed for activated mouse T cells, and further demonstrated combinational workflows in which retroviral transduction can be applied post-EP to enable introduction of biosensors or genetic rescue constructs. For BMDMs and HSCs, we evaluated multiple pre-programmed EP conditions and identified high-performing programs that support robust RNP delivery and efficient gene knockout. Across all cell types, these optimized workflows achieved high electroporation efficiency, cell viability, and reproducibility, with minimal disruption to proliferation, phenotype or the ability to differentiate. Together, these findings demonstrate that MaxCyte ExpERT electroporation is a clinically relevant, scalable, and functionally robust platform for genome editing in primary mouse hematopoietic lineage cells.

## Materials and Methods

| Reagents | Source | Identifier |
| --- | --- | --- |
| Alt-R CRISPR-Cas9 crRNA | IDT | N/A |
| Alt-R® CRISPR-Cas9 tracrRNA, ATTO™ 550 | IDT | Cat# 1075928 |
| APC Anti-CD34; Clone MEC14.7 | BioLegend | Cat# 119310 |
| Pacific Blue Anti-CD38; Clone 90 | BioLegend | Cat# 102719 |
| PerCP-Cy5.5 Anti-CD90.2; Clone 53-2.1 | BioLegend | Cat# 140321 |
| Super Bright 702 Anti-F4/80; Clone BM8 | eBioscience | Cat# 67-4801-82 |
| Cas9-NLS purified protein | UC Berkeley QB3<br>MacroLab | N/A |
| CleanCap EGFP mRNA | TriLink | Cat# L-7201 |
| DPBS (1X) | Corning | Cat# 21-031-CV |
| Easysep™ Mouse CD8<sup>+</sup> T Cell Isolation Kit | STEMCELL<br>Technologies | Cat# 19853 |
| EasySep™ Mouse Hematopoietic Progenitor Cell Isolation Kit | STEMCELL<br>Technologies | Cat# 19856A |
| Fixable Viability Dye eFluor 780 | eBioscience | Cat# 65-0865-18 |
| HyClone HyPure Water, Cell Culture Grade | Cytiva | Cat# SH30529.LS |
| HyClone™ Water, Molecular Biology Grade | Cytiva | Cat# SH3053802 |
| InVivoMAb anti-mouse CD28 | BioXCell | Cat# BE0015-1 |
| InVivoMAb anti-mouse CD3 | BioXCell | Cat# BE0002 |
| MaxCyte® Electroporation Buffer | MaxCyte | Cat# EPB-1 |
| Recombinant human IL-2 | PeptoTech | Cat# 200-02 |
| Recombinant murine SCF | Peptotech | Cat# 250-03 |
| Recombinant murine IL-3 | Peptotech | Cat# 213-13 |
| Recombinant murine IL-6 | Peptotech | Cat# 216-16 |
| RPMI 1640 | Corning | Cat# 10-040-CV |
| Iscove's Modification of DMEM (IMDM) | Corning | Cat# CA45000-366 |
| Trizma® hydrochloride solution (10X) | Sigma | Cat# T2819-1L |
| Trypan Blue Stain | Invitrogen | Cat# 15250061 |
| **Equipment** | **Source** | **Identifier** |
| 0.2 ml PCR tube strips | VWR | Cat# 53509-304 |
| 6 well plates, TC treated | VWR | Cat# 10062-892 |
| 6 well plates, non-TC treated | VWR | Cat# 10861-554 |
| Cell strainer 40 µM | VWR | Cat# 732-2757 |
| Cell strainer 70 µM | VWR | Cat# 76327-100 |
| Cell lifter | Fisher brand | Cat# 08100240 |
| BD FACSymphony™ A1 Cell Analyzer | BD Biosciences | N/A |
| EasySep Magnet | STEMCELL Technologies | Cat# 18000 |
| Eppendorf 5910 R Centrifuge | Eppendorf | Cat# 2231000623 |
| ExPERT ATx™ | MaxCyte | N/A |
| ExPERT™ Docking Station | MaxCyte | Cat# DKS01-01 |
| MaxCyte® Processing Assembly_OC-25x3™ | MaxCyte | Cat# SOC-25x3 |
| MaxCyte® Processing Assembly_R-50x3™ | MaxCyte | Cat# ER050U8-03 |
| MaxCyte® Processing Assembly_R-50x8™ | MaxCyte | Cat# ER050U8-03 |
| MiniAmp Plus Thermal Cycler | Thermo | Cat# A37029 |
| Px5™ Workflow Rack | MaxCyte | Cat# RKPA5-02 |
| Px6s™ Strip Workflow Rack | MaxCyte | Cat# RKS06-02 |

### Detailed procedures

C57BL/6-Tg(TcraTcrb)1100Mjb/J (OT-I) mice were used for primary CD8⁺ T cell activation experiments, with spleen and lymph nodes harvested for cell isolation. Primary bone marrow– derived monocytes/macrophages (BMDMs) and hematopoietic stem cells (HSCs) were isolated from femurs and tibias of C57BL/6J wild-type (WT) mice. All founder mice were obtained from The Jackson Laboratory and bred at the Centre for Molecular Medicine and Therapeutics (CMMT) Animal Facility in accordance with institutional guidelines (approval protocols #A23-0194 and A23-0282).

### Section1: Procedures for mouse primary CD8^+^ T cells

#### Day 0 Mouse primary CD8^+^ T cell isolation and culture

*Timing: 3.5 hours*

1. Harvest spleen and lymph nodes (Inguinal, axillary, brachial, cervical nodes, ensuring there is no remaining fat tissue attached) from OT-I mice, and place into 4 mL of ice-cold wash buffer (RPMI 1640 with 2% fetal bovine serum (FBS)). *NOTE: All FBS used in this manuscript was heat inactivated at 55°C for 30 minutes*.
2. Place a sterile 70 µm cell strainer on a 50 mL conical tube and place spleen and lymph nodes on top. Dissociate tissues using the back of the sterile plunger of a 1 mL or 3mL syringe through mashing. *NOTE: Tissues should be processed immediately after collection to preserve cell viability. If processing is delayed, the single-cell suspension may be maintained in ice-cold wash buffer on ice*.
3. Isolate CD8^+^ T cells from the spleen and lymph nodes using negative selection with an EasySep™ mouse CD8^+^ T cell isolation kit following the manufacturer’s protocol.
4. After isolation, take a 10 µL aliquot and dilute in 90 µL of trypan blue to count using a hemocytometer. *NOTE: To ensure accurate cell counts, automated cell counters are not recommended unless validated using trypan blue–stained samples*.
5. Centrifuge CD8^+^ T cell suspension at 400×g for 5 minutes, aspirate the supernatant, and resuspend in T-cell medium (TCM) at a 2×concentrated density of 3×10^6^ cells/mL. *NOTE: TCM was prepared by adding 10% FBS, 1% penicillin/streptomycin, 2 mM L-glutamine, and 50 µM* β*-mercaptoethanol in RPMI 1640 medium*.
6. Prepare CD8^+^ T-cell activation plate and medium:

- Make coating buffer by diluting 10×Trizma solution in cell culture grade water to achieve 1× Trizma solution and add InVivoMAb anti-mouse CD3 to final concentration of 5 µg/mL.
- Add 2 mL of InVivoMAb anti-mouse CD3 containing coating buffer per well in a 6-well cell culture plate. *NOTE: Select a suitable culture vessel appropriate for the final culture volume and surface area to maintain a consistent anti-mouse CD3 coating density of 1.05 µg/cm*^2^.
- Incubate the plate at 37°C for at least 2 hours.
- Prepare T-cell activation medium by adding 2× concentrated InVivoMAb anti-mouse CD28 (1.0 µg/mL) and 2× concentrated rhIL-2 (200 U/mL) in TCM.
- Aspirate anti-mouse CD3 solution and wash the cell culture plate with 1 mL of 1×DPBS twice. *Note: Take care to not let wells become dry in between and after the washes*.
7. Add 1:1 (v/v) ratio of prepared T-cell activation medium and isolated CD8^+^ T cells suspension. *NOTE: The final cell density is 1.5*×*10*^6^ *cells/mL and cells are plated at a surface density of 1.27×10^6^ cells/cm*^2^ *in a 6-well cell culture plate. Final concentrations of soluble anti-mouse CD28 and rhIL-2 are 0.5 µg/mL and 100 U/mL, respectively*.
8. Culture the CD8^+^ T cells at 37℃ in a humidified incubator with 5% CO_2_ for 24 – 48 hours.

#### Day 1 or 2 CRISPR-Cas9 Ribonucleoprotein (RNP) and mRNA electroporation

*Timing: 2 hours*

*NOTE: Electroporation can be conducted 1 or 2 days post-activation depending on the experimental needs. CD8^+^ T cells activated using this protocol do not divide before 36 hours*^39^*[dough*.

1. To generate sgRNA, mix 200 µM crRNA and 200 µM µM tracrRNA (ATTO-550) at a 1:1 ratio. *NOTE: Lyophilized crRNA and tracrRNA were resuspended in nuclease-free water to a final stock concentration of 200 µM (200 pmol/µL)*.
2. Incubate at 95℃ for 5 minutes using a thermal cycler.
3. Cool the sgRNA for 15 minutes at ambient temperature in the dark.
4. Combine 25 µL of 100 µM sgRNA and 20 µL of 40 µM Cas9 for a final RNP complex concentration of 22.2 µM (2:1 sgRNA:Cas9 ratio).
5. Incubate the RNP complex for 10 minutes at RT.
6. Keep the RNP complex in the dark until use.
7. Thaw eGFP mRNA to use as a control on ice.
8. For each sample with RNP, pipet 4.5 µL of 22.2 µM RNP complex into a sterile 0.2 mL PCR tube. The final concentration will be 2 µM (2 pmol/µL) of RNP complex in 50 µL of the total electroporation reaction volume.
9. For each sample with eGFP mRNA, pipet 5 µL of 1 µg/µL mRNA into a sterile and nuclease-free 0.2-mL PCR tube. The final concentration will be 100 µg/mL of mRNA in 50 µL of the total electroporation reaction volume.
10. Pre-warm 3 mL TCM containing 100 U/ml rhIL-2 per electroporation condition in a 6-well plate in a 37℃ incubator.
11. Collect activated mouse CD8^+^ T cells from the activation plates by thorough pipetting to detach cells and visually inspecting the wells under a microscope to confirm maximal cell recovery.
12. Count cells and measure viability using trypan blue. NOTE: Ensure that cells have healthy morphology and that cell viability is high. Starting cell health is very important to a successful electroporation. T cell viability should be at least 80% or greater, ideally 90% or higher.
13. Calculate the number of cells needed for the experiment.

- Use Table 2 to calculate number of cells based on the desired electroporation volume using a final concentration of 1×10^8^ cells/mL.
- Increase the starting total cell number by 20% to account for pipetting losses (see Table 2 row E). *NOTE: The final cell concentration should be 1*×*10*^8^ *cells/mL or greater. If there are not enough cells, consider lowering the electroporation volume first, in combination with the appropriate electroporation cuvette (“Processing Assembly”). (e.g., 2*×*10*^6^ *cells in 20 µL). The standard protocol uses 5*×*10*^6^ *cells in 50 µL for consistency and ease of calculation*.
14. Transfer the desired number of cells into a 15 mL or 50 mL conical tube.
15. Centrifuge cells at 200×g for 5 minutes with low brake (set deceleration setting on centrifuge to 30% of maximum depending on the model).
16. Aspirate the supernatant and resuspend the cell pellet with MaxCyte electroporation buffer. Use a volume that is at least 5 times that of the pellet to wash thoroughly.
17. Centrifuge cells at 200×g for 5 minutes with low brake.
18. Aspirate supernatant and resuspend cell pellet in MaxCyte electroporation buffer such that the volume including the pellet is the target resuspension volume.

- First, add MaxCyte electroporation buffer to reach half of the target resuspension volume.
- Gently resuspend the pellet by raking the tube across a grill of a biosafety cabinet or by tapping.
- Measure the exact volume with a micropipette.
- Calculate the remaining volume needed to reach the target resuspension volume.
19. For each sample, perform the following steps in full before continuing to the next sample:

- Add 45 µL of concentrated cell stock to the tube with 5 µL of loading agents (buffer only or RNP or eGFP mRNA).
- Mix by gently pipetting 4 - 5 times to avoid bubble formation.
- Take the entire 25 µL or 50 µL volume to add to one empty well of the OC-25×3 or R-50×8 Processing Assembly, respectively.
- Prepare docking station for R-50×8 Processing Assembly.
20. Electroporate with the “Expanded T Cell 3” or “Expanded T Cell 4” protocol for OT-1 T cells.
21. For each sample, transfer cells from the Processing Assembly into 3 mL of pre-warmed TCM in one well of a 6-well plate (final concentration of 1.5 × 10^6^ cells/mL).

- Letting cells sit undisturbed for 5 – 15 minutes post-EP is beneficial before replating in TCM.
22. Add 50 µL of TCM to wash the well of the Processing Assembly, pipetting 1-2 times to wash out any remaining cells, and add to corresponding well in the plate.
23. Distribute cells evenly in the cell culture plate by swirling and horizontal rocking.
24. Incubate overnight in a cell culture incubator (37°C, 5% CO_2_, humidity-controlled).

**Table 1.**
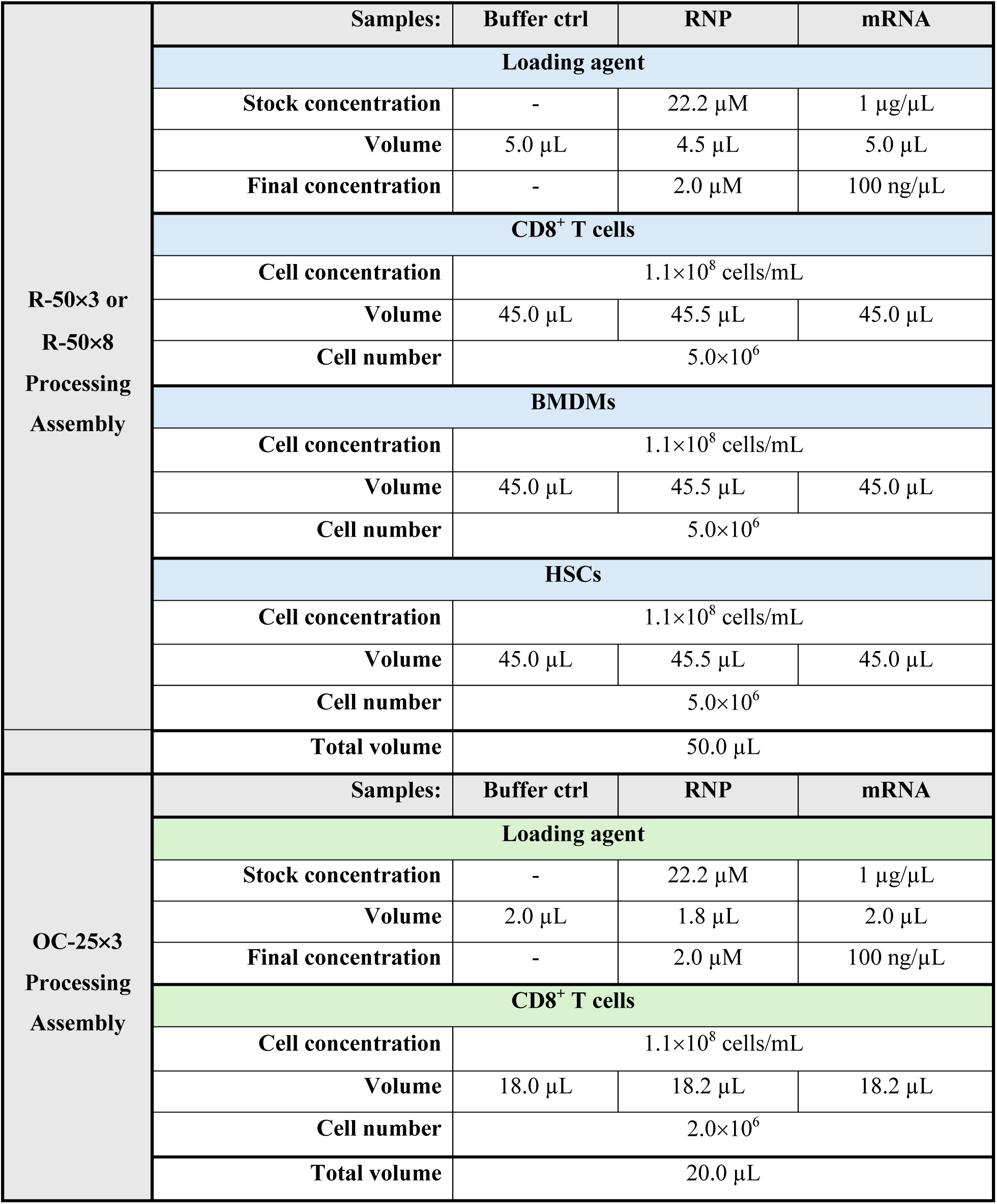
Calculation for loading agents and electroporation reaction volume.

**Table 2.** Cell number calculation applicable to across all cell types used in this study.

| A | Parameter | Formula | Example |
| --- | --- | --- | --- |
| B | # of samples | N | 8 |
| C | Electroporation volume | V | 50.0 μL (0.050 mL) |
| D | Min. # of cells required | $\text{Total}_{\min} = (N+2) \times 1.0 \times 10^8 \text{ cells/mL} \times V$ | $\text{Total}_{\min} = 5.0 \times 10^7 \text{ cells}$ |
| E | Total cells to collect | $\text{Total}_{\min} \times 120\%$ | $6.0 \times 10^7 \text{ cells}$ |
| F | Resuspension volume | $(\text{Total}_{\min} \times 120\%) \div 1.1 \times 10^8 \text{ cells/mL}$ | 545 μL |

#### Day 2 or 3 (1 day post-electroporation)

*Timing: 1.5 hours*

1. Count cells and measure viability.
2. Check that cells still have healthy morphology and high cell viability.
3. Check EP efficiency with flow cytometry:

- Take 50,000 - 100,000 cells per group for flow cytometry.
- Measure cell viability by live/dead stain and check for fluorescence in expected channel (ex. ATTO-550 from tracrRNA in PE or its equivalent channel).
4. For each sample, replate T cells in TCM at 1.5×10^6^ cells/mL in a 6-well plate.

**Day 4 – 5** Continue expansion and culture of CD8⁺ T cells for functional or phenotyping experiments. In this study, transient culturing in low-glucose media was used as an example application for phenotyping^40^.

### Section 2: Procedures for mouse primary BMDMs

#### Day 0 Mouse primary BMDMs isolation and electroporation

1. Collect femur and tibia from WT mice and keep the bones in 1 mL of ice-cold wash buffer (RPMI 1640 + 2% FBS).
2. Isolate the bone marrow from the femur and tibia by cutting both ends of the bones with sterile scissors. Next, use an 18 gauge needle to puncture a hole at the bottom of a 0.5 mL Eppendorf tube and place the bones inside. Put the 0.5 mL Eppendorf inside a 1.5 mL Eppendorf tube and centrifuge at 3300×g for 15 seconds to flush the bone marrow into the 1.5 mL Eppendorf tube.
3. Remove the 0.5 mL tube containing the flushed bones and discard. Resuspend the bone marrow pellet in 1 mL RBC lysis buffer and incubate for 5 minutes at RT. In the meantime, prepare one 50 mL tube per mouse and place a 40 µm cell strainer to each of them. Pre-wet the cell strainers with by adding 10 mL of macrophage media (Mac media) through the strainer and into the 50 mL tube. *NOTE: Mac media was prepared by adding 10% FBS, 1% penicillin/streptomycin in RPMI 1640 medium*.
4. Transfer the cells in lysis buffer into a 50 mL tube containing Mac media through a 40 um pre-wet cell strainer, effectively stopping the lysis reaction.
5. Wash the RBC lysis buffer out by centrifuging the cells at 400×g for 5 minutes at RT.
6. Resuspend pellet in Mac media for counting.
7. Prepare sgRNA and RNP complex as described in Section 1, Day 1, Steps 1 - 9.
8. Wash, prepare and electroporate cells using the calculations in Table 2 starting at Step 14 using “Optimization 0-2 (Opt 0-2)”, “Hematopoietic stem cell-4 (HSC-4)”, or “Hematopoietic stem cell-6 (HSC-6)” protocols installed on the electroporator.

#### Day 1 post-electroporation

1. Initiate macrophage differentiation, according to the previous protocol^41^.
2. For each sample, resuspend electroporated cells in Mac differentiation media for replating. *NOTE: Mac differentiation media was prepared by adding 10% FBS, 1% penicillin/streptomycin, 1% HEPES, and 20% L929 supernatant media in RPMI 1640 medium, as previously described*^41^*. Alternatively M-CSF could be used in place of L929 supernatant*^42^.
3. Replate cells as 2×10^6^ cells/mLin a 6-well cell culture plate or as 7×10^6^ cells/mL in a 10 cm TC dish for differentiation for 7 days. Replace 50% of the culture medium on days 3 and 5 after plating.

### Section 3: Procedures for mouse primary HSCs

#### Day 0 Mouse primary HSCs isolation and culture

*Timing: 3.5 hours*

1. Harvest femur and tibia from WT mice as in Section 2, Day 0, Steps 1 – 3.
2. Isolate HSCs from the bone marrow by using EasySep™ mouse hematopoietic progenitor cell isolation kit following the manufacturer’s protocol.
3. After isolation, take a 10 µL aliquot and dilute in 40 µL of trypan blue to count.
4. Centrifuge isolated HSCs suspension at 400×g for 5 minutes at RT, aspirate the supernatant, and resuspend in complete IMDM(cIMDM) at a density of 1.0×10^6^ cells/mL *NOTE: cIMDM was prepared by adding 20% FBS, 1% penicillin/streptomycin, IL-3 (20 ng/mL), IL-6 (30 ng/mL), and SCF (50 ng/mL)*.
5. Plate the HSCs in a 6-well non-TC treated plate and culture at 37℃ in a humidified incubator with 5% CO_2_ for 24 hours.

#### Day 1 HSCs expansion

*Timing: 30 minutes*

1. Double the culture media volume by adding cIMDM.

#### Day 2 Continue with HSCs expansion

*Timing: 1 hour*

1. Harvest HSCs from the culture plates by vigorous pipetting to collect maximal number of cells.
2. Take a 10 µL aliquot and dilute in 40 µL of trypan blue to count.
3. For each sample, replate HSCs in cIMDM at a density of 1.0×10^6^ cells/mL in a 6-well non-TC treated plate.

#### Day 3 eGFP mRNA and RNP electroporation

*Timing: 2 hours*

1. Prepare sgRNA and RNP complex as described in Section 1, Day 1, Steps 1 - 9.
2. Collect HSCs from the 6-well non-TC treated plates.
3. Count, prepare, and electroporate cells using the calculations in Table 2 with one of the following protocols: “Opt 0-2”, “HSC-4”, or “HSC-6”. *NOTE: Refer to Section 1, Steps 12-24 for details, and the electroporation programs used for HSCs are the same as those for mouse primary BMDMs*.
4. Once the electroporation is completed, transfer cells from the Processing Assembly into 5 mL of pre-warmed complete IMDM in one well of a 6-well non-TC treated plate to make a final concentration of 1.0×10^6^ cells/mL.

#### Day 4 (1 day post-electroporation)

*Timing: 1.5 hours*

1. Count and resuspend the electroporated HSCs in cIMDM at 1.0×10^6^ cells/mL in a 6-well non-TC treated plate. *NOTE: If positive control included, take 50,000 – 100,000 cells for flow cytometry as indicated in Section 1, Day 2 or 3, Step 3*.

#### Day 5

The electroporated HSCs can be used for downstream experiments, including colony assay and BM transplantation in irradiated mice^43,44^.

## Results

### Optimized electroporation of OT-I T cells enables high-efficiency mRNA and RNP delivery without compromising viability

When working with primary cells, factors such as cell viability and transfection efficiency are essential for establishing a reliable workflow for gene editing^45^. OT-I TCR-transgenic (Tg) mice are widely used to investigate how CD8^+^ T cells respond to their cognate antigens^46^. In this study, we used OT-I T cells that were 24 hours (Day 1) and 48 hours (Day 2) post-initial activation. Electroporation of OT-I T cells was performed using either the “Expanded T cell 3” (Exp T3, low energy) or “Expanded T cell 4” (Exp T4, high energy) protocol. As cargo, we delivered either buffer control, eGFP mRNA, or two independent RNP complexes (**Fig. 1a**). Electroporation outcomes were evaluated by flow cytometric analysis of viability and efficiency. Cell viability was assessed 24 hours after the electroporation. Notably, when OT-I cells on Day 1 or Day 2 after activation were electroporated with either the Exp T3 or Exp T4 protocol, more than 95% of cells remained viable, as demonstrated in the representative dot plots (**Fig. 2a, b, Supplementary Fig. 1a**) and quantified in the corresponding bar graphs (**Fig. 2c, d**). These findings indicate that neither protocol compromised T cell viability under the conditions tested. We next quantified the proportion of eGFP-expressing OT-I cells 24 hours after electroporation. Using either the Exp T3 or Exp T4 protocol, transfection efficiency exceeded 95% for both Day 1 and Day 2 OT-I cells, as shown by the representative histograms (**Fig. 2e**) and summarized in the bar graphs (**Fig. 2f**). Day 2 OT-I cells exhibited more than a twofold increase in eGFP geometric mean fluorescence intensity (gMFI) compared to Day 1 cells (**Fig. 2g**). The elevated expression in Day 2 cells is consistent with the continuous increase in translational capacity during the activation of T cells, likely driven by increased mTORC1 signaling at 48 hours post-activation^47^. Exp T3 and Exp T4 exhibited no statistically significant difference in % eGFP^+^, indicating equivalent mRNA delivery efficiency between the two protocols. Lastly, we assessed RNP delivery efficiency. Over 98% of OT-I Day 1 and Day 2 cells were positive for ATTO-550-labeled tracrRNA following electroporation with either RNP1 or RNP2, as shown in the histograms (**Fig. 2h**) and bar graph quantification (**Fig. 2i**). Together, these results demonstrate that both Exp T3 and Exp T4 protocols achieve robust mRNA and RNP delivery into primary OT-I T cells while preserving viability, establishing a reliable and reproducible platform for transient expression or gene-editing applications in primary mouse T cells.

**Fig. 1.**
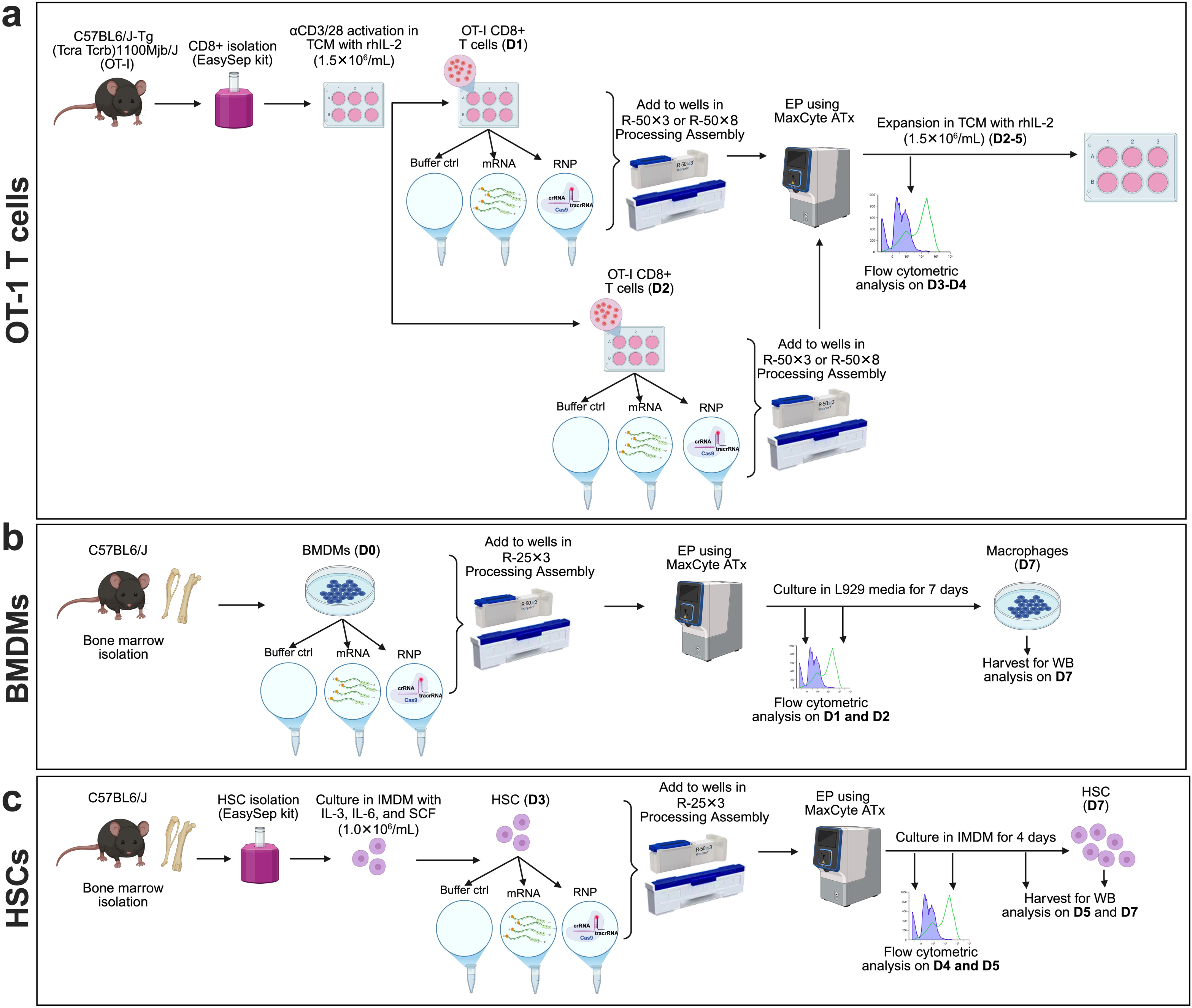
Schematic illustration depicting mouse primary hematopoietic cells and electroporation procedures. **a-c** Schematic illustration depicting OT-I T cell activation/expansion (**a**) bone marrow-derived macrophages (BMDMs) isolation (**b**) hematopoietic stem cells (HSCs) isolation (**c**). Electroporation procedures are shown for each cell type.

**Fig. 2.**
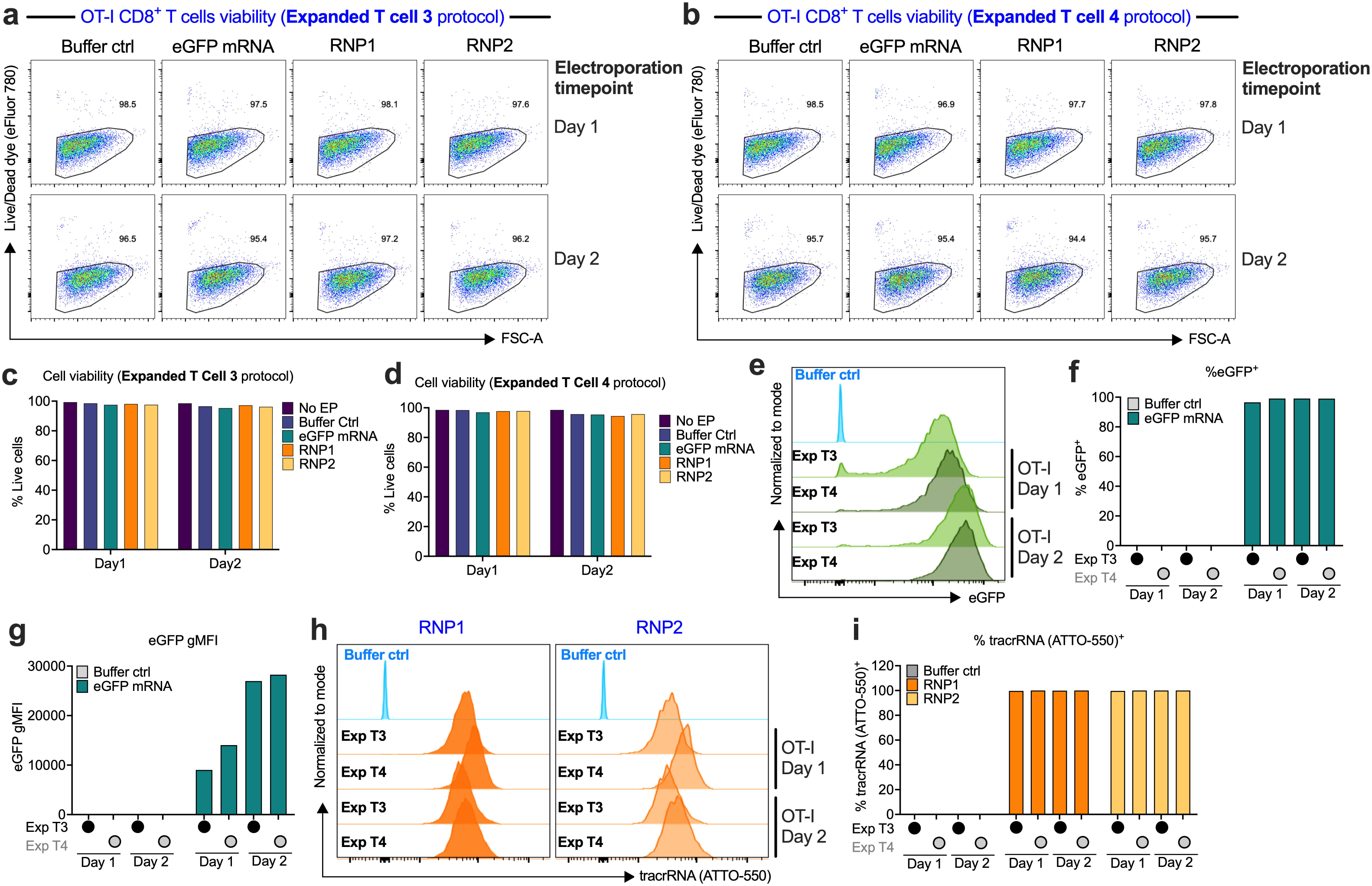
Comparison of Expanded T cell 3 and Expanded T cell 4 electroporation programs demonstrates efficient cargo delivery and high viability in OT-I T cells 24 hr post-electroporation. **a-i** 100,000 OT-I T cells were harvested and assessed by flow cytometry 24 hr post-electroporation. **a, b** Representative flow cytometry plots showing the cell viability following electroporation using Expanded T cell 3 (Exp T3) protocol (**a**) and Expanded T cell 4 (Exp T4) protocol (**b**). Viability was measured using a fixable Live/Dead dye (eFlour 780). **c, d** Quantification of the frequency of viable cells following electroporation using the Exp T3 (**c**) or Exp T4 (**d**) protocols. **e-g** Day 1 or Day 2 activated OT-I T cells were electroporated with eGFP mRNA using either Exp T3 or Exp T4 protocols, and eGFP expression was measured 24 hr post-electroporation. Representative histograms show the distribution of eGFP intensity (**e**). Bar graphs summarize the frequency of eGFP⁺ cells (**f**) and the geometric mean fluorescence intensity (gMFI) of eGFP⁺ cells (**g**). **h, i** Day 1 or Day 2 activated OT-I T cells were electroporated with either RNP1 or RNP2 using Exp T3 or Exp T4 protocols. Representative histograms show the ATTO-550-labeled tracrRNA expression (**h**), and bar graphs quantify the frequency of ATTO-550⁺ cells (**i**).

### Primary OT-I T cells retain growth capacity and show efficient gene-editing following electroporation

Following electroporation, we cultured OT-I T cells and monitored their expansion while confirming gene-editing efficiency by Western blotting (**Fig. 3a**). Under the Exp T3 protocol, Day 1 cells displayed reduced early expansion but nonetheless increased cell number by approximately 3-fold over 72 hours (**Fig. 3b**), whereas Day 2 cells expanded around 4-fold during the same period (**Fig. 3c**). By the end of culture, fold expansion was only minimally different from buffer control T cells in both Day 1- and Day 2-electroporated groups (**Fig. 3b, c**). Comparable results were observed with the Exp T4 protocol (**Fig. 3d, e**); however, expansion of Day 2 cells was noticeably lower than that of Day 2 cells electroporated with Exp T3, suggesting that the expansion of more activated, proliferating T cells may be more sensitive to the higher-energy Exp T4 protocol.

**Fig. 3.**
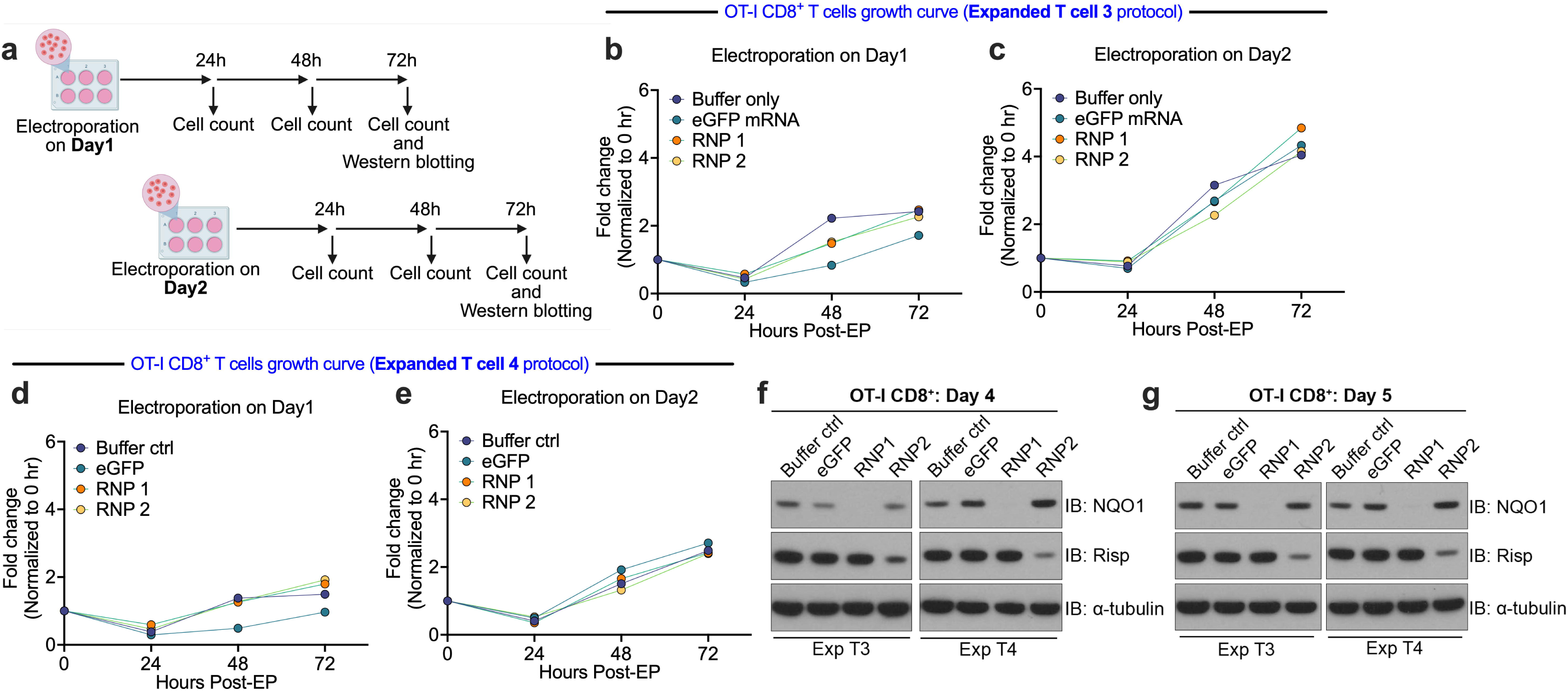
Electroporated OT-I T cells retain proliferative capacity and support efficient gene knockout. **a** Schematic of the experimental workflow used to quantify cell expansion following electroporation. **b, c** Day 1 (**b**) or Day 2 (**c**) activated OT-I T cells were cultured for 72 hr following electroporation using the Expanded T cell 3 protocol. Relative proliferation was assessed by cell counting at 24 hr intervals. **d, e** Day 1 or Day 2 activated OT-I T cells were cultured for 72 hr following electroporation using the Expanded T cell 4 protocol. Relative proliferation was assessed by cell counting at 24 hr intervals. **f** Following electroporation, OT-I T cells were harvested and analyzed by immunoblotting for NQO1 or Risp expression. α-tubulin was used as a loading control.

OT-I cells electroporated on Day 1 were expanded for 3 days, after which cell pellets were collected and lysed for Western blot analysis. A protein normally upregulated by the gene-product targeted by RNP1 was efficiently knocked out using both the Exp T3 and Exp T4 protocols (**Fig. 3f, top blots**). In contrast, the protein encoded by the gene targeted by RNP2 showed only a modest reduction in expression (**Fig. 3f, middle blots**). Given that RNP2 electroporation efficiency exceeded 95% (**Fig. 2h, i**), the limited knockout likely reflects suboptimal sgRNA performance rather than delivery inefficiency, highlighting the importance of testing multiple independent gRNA to ensure efficient gene editing and loss of protein expression. Similar patterns were observed in Day 2 OT-I cells, with robust knockout of the RNP1 target and minimal reduction of the RNP2 target across both electroporation protocols (**Fig. 3f**), indicating that gene editing outcomes were consistent regardless of activation state at the time of electroporation. Together, these results show that electroporated OT-I T cells retain their doubling capacity across protocols and activation states at the time of electroporation, with only minor sensitivity of Day 2 cells to the higher-energy Exp T4 condition. Moreover, robust knockout of the RNP 1 target demonstrates that both protocols support efficient gene editing, while the limited effect of RNP 2 highlights sgRNA quality as the principal determinant of editing outcomes using these protocols. These data also highlight the need for an efficient, reproducible transfection method in order to avoid uncertainty with whether low knock-out efficiency is due to a target sequence issue or transfection issue.

### Electroporation does not exacerbate ER stress–related mRNA expression in low-glucose CD8⁺ T cells

Glucose availability is critical for N-linked protein glycosylation and limiting glucose can provoke ER stress. In addition, electroporation has also been reported to trigger ER stress^48^. To assess whether reduced glucose and/or electroporation alter the nutrient-mediated ER stress response in CD8⁺ T cells, we measured mRNA expression of *Atf4* and *Ddit3* (encoding Chop), canonical markers of nutrient and ER stress, in CD8⁺ T cells under nutrient-restricted conditions^49^. Both transcripts were consistently upregulated under low-glucose conditions (**Supplementary Fig. 2a, b**). However, electroporation did not further increase *Atf4* or *Ddit3* mRNA expression, indicating that this potentially stress-inducing procedure does not exacerbate ER stress in primary CD8⁺ T cells. These findings demonstrate that while low-glucose CD8⁺ T cells mount a stress-responsive transcriptional program, electroporation does not amplify ER stress–related mRNA expression, underscoring the robustness of this workflow for functional studies under metabolic perturbation.

### Sequential electroporation and retroviral transduction enable efficient co-expression of CRISPR-Cas9 RNP and a reporter construct in OT-I T cells

Gene knockout studies are frequently complemented by gene rescue experiments to validate the specific functional contribution of target genes. However, knock-in approaches often exhibit limited efficiency and typically require isolation of single integration clones for downstream analysis, posing a barrier for short-lived *in vivo* mouse T cell studies. To establish knockout-rescue workflow, we tested a sequential platform combining electroporation-mediated RNP delivery (knockout) with retroviral gene expression (rescue) in activated OT-I T cells. Day 1 activated OT-I T cells were electroporated using the Exp T3 protocol to deliver RNP complexes. Following a 24 hr recovery period, cells were retrovirally transduced on Day 2 by spin-fection with a GFP reporter construct and expanded for an additional 48 hr (**Supplementary Fig. 3a**). Flow cytometric analysis revealed robust co-expression of GFP and ATTO-550–labeled tracrRNA, indicating successful sequential delivery of viral and non-viral cargos within the same cell population (**Supplementary Fig. 3b**). The majority of GFP⁺ cells retained detectable ATTO-550–labeled tracrRNA fluorescence, demonstrating that electroporation does not impair subsequent retroviral transduction efficiency and that RNP delivery and editing can proceed alongside viral transduction and cell expansion. These findings establish a modular workflow in which rapid, non-viral genome editing can be coupled with optional viral delivery of reporter or rescue constructs, enabling flexible gene perturbation and functional complementation strategies in primary T cells while minimizing the downstream need for single clone isolation steps often used with knock-in strategies.

### High-efficiency electroporation preserves viability and differentiation of mouse primary BMDMs

Given the robust gene-editing efficiency achieved in mouse primary OT-I T cells using the MaxCyte ExPERT platform, we next assessed its performance in additional primary cell types known to be exquisitely sensitive and difficult to genetically modify, including BMDMs and HSCs^50,51^. Bone marrow was harvested from C57BL/6J mice and electroporated using pre-programmed protocols “Opt 0–2”, “HSC-4”, and “HSC-6”, followed by differentiation into BMDMs over 7 days (**Fig. 1b**). Viability was assessed 24 and 48 hours after electroporation. Across buffer control (ctrl), eGFP mRNA, and RNP cargo conditions, BMDMs displayed 88.7– 94.4% viability, as shown in representative dot plots (**Fig. 4a, b; Supplementary Fig. 1b**). Quantification confirmed no significant difference between electroporated groups and either non-electroporated or buffer-only ctrl at both time points (**Fig. 4c, d**). These results demonstrate that BMDMs maintain high viability following electroporation, underscoring the suitability of this platform for gene-editing applications in difficult to genetically modify primary myeloid lineages.

**Fig. 4.**
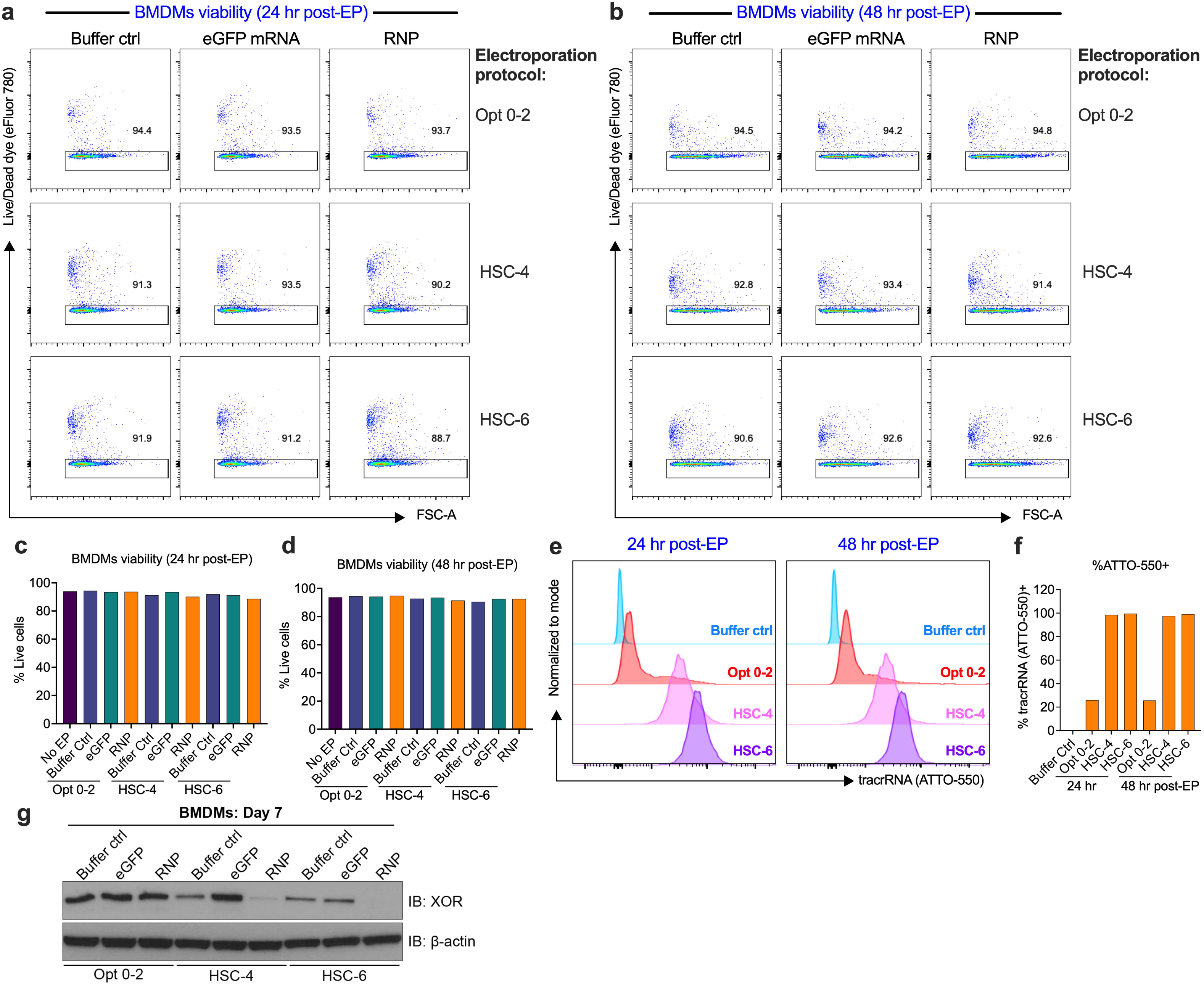
Optimized electroporation supports efficient RNP delivery and gene editing in primary BMDMs. Isolated bone marrow cells were electroporated with buffer control (ctrl), eGFP mRNA, or RNP using three electroporation protocols: Opt 0–2, HSC-4, and HSC-6. **a, b** Representative flow cytometry dot plots showing the frequency of viable cells at 24 hr (**a**) and 48 hr (**b**) post-electroporation. **c, d** Quantification of viable cell frequencies in non-electroporated (No EP) and electroporated (EP) groups at 24 hr (**c**) and 48 hr (**d**) post-electroporation. **e** Representative histograms showing electroporation efficiency based on ATTO-550–labeled tracrRNA fluorescence at 24 hr and 48 hr post-electroporation. **f** Quantification of ATTO-550⁺ cells at 24 hr and 48 hr post-electroporation. **g** Following electroporation, bone marrow cells were differentiated into macrophages for 7 days as described in the Methods and analyzed by immunoblotting for XOR expression. β-actin was used as a loading control.

To further evaluate the platform performance, we examined cargo delivery and gene-editing performance in BMDMs. eGFP expression was not detected following electroporation (**Supplementary Fig. 3a**), which may reflect intrinsic RNase activity or reduced translational capacity in macrophages^52,53^. To assess RNP delivery efficiency, we measured uptake of ATTO-550-labeled tracrRNA by flow cytometry. Approximately 20% of BMDMs electroporated with the Opt 0–2 protocol was ATTO-550 positive at both 24 and 48 hours, whereas the HSC-4 and HSC-6 protocols yielded > 95% ATTO-550⁺ cells (**Fig. 4e, f**). Among these, HSC-6 provided the most robust performance as indicated by the highest ATTO-550 signal. Consistent with the observed delivery efficiency, Western blot analysis confirmed that RNP electroporation with the HSC-6 protocol effectively knocked out the target protein in BMDMs (**Fig. 4g**). Importantly, electroporation with the HSC-6 protocol did not impair macrophage differentiation, as Day 7 BMDMs showed comparable F4/80 expression across all conditions (**Supplementary Fig. 3b**). Together, these results identify HSC-6 as an optimal electroporation protocol for efficient and reproducible RNP delivery and gene editing in mouse primary macrophages.

### Optimized electroporation supports high-efficiency gene editing in mouse primary HSCs

Having confirmed efficient delivery in macrophages, we next tested HSCs, a distinct BM-resident cell type used for therapeutic genome engineering^22^. HSCs were isolated from C57BL/6J bone marrow and expanded for 3 days prior to electroporation. Day 3 HSCs were electroporated using the pre-programmed protocols “Opt 0–2”, “HSC-4”, and “HSC-6” (**Fig. 1c**). Viability remained consistently above 90% at 24 hours post-electroporation and was similarly maintained at 48 hours across all cargo conditions, including buffer ctrl, eGFP mRNA, and RNP complex (**Fig. 5a-d**). Similar to BMDMs, eGFP expression was not detected following electroporation in HSCs (**Supplementary Fig. 3c**). To assess RNP delivery efficiency, we measured uptake of ATTO-550-labeled tracrRNA by flow cytometry. Approximately 80% of HSCs electroporated with Opt 0–2 or HSC-4 were ATTO-550⁺ at 24 hours, with a modest decrease of 5–10% observed at 48 hours post-electroporation (**Fig. 5e, f**). By contrast, HSC-6 yielded near-uniform delivery, with ∼99% ATTO-550⁺ cells maintained stably at both 24 and 48 hours (**Fig. 5e, f**). Cells were expanded until day 5 and day 7, and pellets were collected for Western blot analysis. Consistent with the delivery results, HSC-6 demonstrated the most robust gene knockout, an effect that was preserved in day 7 cells (**Fig. 5g**).

**Fig. 5.**
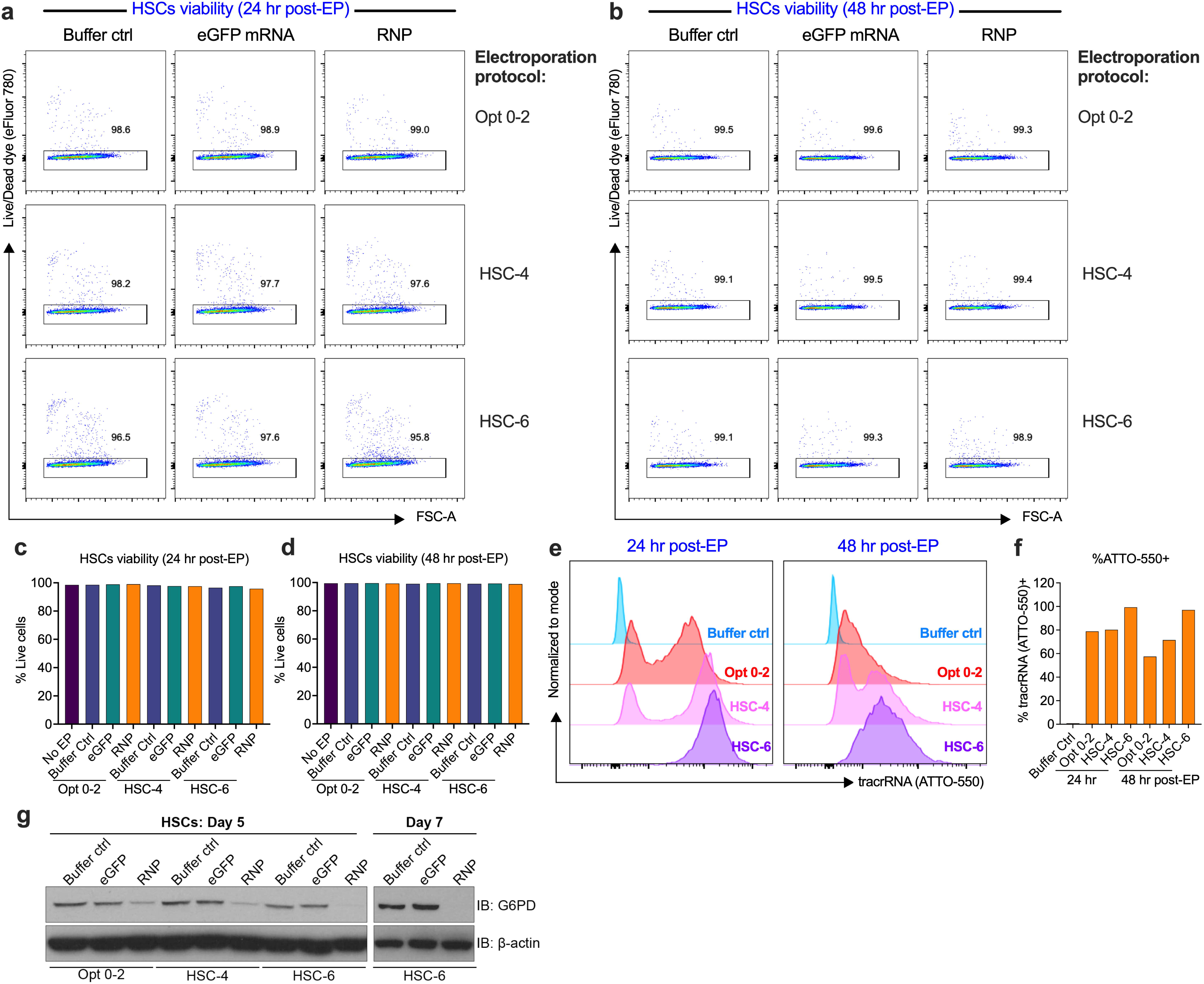
Optimized electroporation achieves efficient gene editing in primary mouse HSCs. HSCs were electroporated with buffer control (ctrl), eGFP mRNA, or RNP using three electroporation protocols: Opt 0–2, HSC-4, and HSC-6. **a, b** Representative flow cytometry dot plots showing the frequency of viable cells at 24 hr (**a**) and 48 hr (**b**) post-electroporation. **c, d** Quantification of viable cell frequencies in non-electroporated (No EP) and electroporated (EP) groups across the three protocols at 24 hr (**c**) and 48 hr (**d**) post-electroporation. **e** Representative histograms showing RNP delivery efficiency based on ATTO-550–labeled tracrRNA fluorescence at 24 hr and 48 hr post-electroporation. **f** Quantification of ATTO-550⁺ cells in buffer ctrl and EP groups at 24 hr and 48 hr post-electroporation. **g** Immunoblot analysis of HSCs harvested on Days 5 and 7 post-electroporation assessing G6PD protein expression.

During extended *in vitro* culture, a proportion of HSCs undergo spontaneous differentiation by Day 7^54^. To control for this effect and focus analysis on the stem/progenitor compartment, cells were first gated on CD34⁺ populations to enrich for short-term HSCs, followed by sub-gating based on CD38 and CD90.2 expression. No significant differences were observed in the frequencies of CD38⁺ or CD90.2⁺ cells within the CD34⁺ compartment between non-electroporated (No EP) and electroporated (EP; HSC-6 protocol) conditions, indicating that electroporation does not promote spontaneous HSC differentiation and preserves key stem-cell phenotypic markers (**Supplementary Fig. 3d**). Together, these findings identify HSC-6 as the optimal protocol for efficient and durable RNP-mediated gene editing in primary mouse HSCs.

## Discussion

Primary mouse hematopoietic cells are foundational to both mechanistic immunology and translational cell engineering, yet their genetic manipulation remains heavily dependent on viral delivery platforms that are costly, time-intensive, and biologically confounding due to genomic integration, innate sensing of exogenous nucleic acid, and prolonged transgene expression^55^.

Non-viral approaches provide a scalable alternative, but in sensitive primary cells, achieving efficient delivery without perturbing phenotype or function requires careful optimization. In this study, we address this bottleneck by optimizing electroporation workflows using the MaxCyte ExPERT platform across three distinct primary mouse hematopoietic lineages, CD8⁺ T cells, BMDMs, HSCs, demonstrating robust delivery while preserving viability, differentiation capacity, and functionality. To our knowledge, this is the first study and comprehensive workflow for highly efficient gene editing across these primary mouse hematopoietic lineages.

In activated OT-I T cells, both the Exp T3 and Exp T4 programs supported high-efficiency mRNA and RNP delivery with minimal impact on viability, establishing electroporation as a reliable platform for transient expression and genome editing in primary T cells. Notably, Exp T3 and Exp T4 yielded equivalent delivery efficiency, but subtle differences in proliferative capacity emerged under the higher-energy Exp T4 condition. These findings suggest a practical balance between delivery energy and proliferative robustness that may depend on activation state rather than intrinsic delivery efficiency. Because starting cell density and viability were carefully controlled prior to electroporation, these differences are unlikely to reflect confounding input variability. From an operational perspective, Exp T3 therefore represents a suitable default when preservation of proliferative capacity is critical, whereas Exp T4 may be preferable for experiment-dependent applications requiring higher energy with acceptable reductions in cell growth.

Electroporation is frequently assumed to impose substantial physiological stress on primary cells^56^, raising concern that it may confound downstream readouts, particularly in metabolic or stress-response studies central to the field of immunometabolism. Consistent with this concern, both nutrient limitation and electroporation have been reported to activate ER stress pathways in other systems^48,49^. Here, we demonstrate that although CD8⁺ T cells appropriately induce canonical stress-responsive transcripts such as *Atf4* and *Ddit3* under glucose-limited conditions, electroporation following our workflow does not further induce this response in glucose-replete cultures. These findings establish an important biological guardrail for applying electroporation-based workflows in studies focussing on stress pathways as mechanistic mediators or experimental readouts. Whether higher energy programs such as Exp T4 subtly engage additional stress pathways that could contribute to reduced proliferation remains an important area for future investigation.

In contrast to T cells, eGFP mRNA expression was not detectable in BMDMs and HSCs under the tested conditions, despite robust RNP delivery and efficient gene knockout. Several non-mutually exclusive mechanisms may explain this lineage-specific difference. Myeloid cells are enriched for innate nucleic acid sensing pathways and RNase activity, which may limit stability or translation of exogenous mRNA^52,53^. In addition, translational capacity and expression kinetics may differ across lineages, and the 24–48-hour sampling window used here may not capture transient expression in these cell populations. The successful delivery and functional activity of RNP complexes indicate that cargo entry per se is not the limiting factor. Future optimization strategies may include modulation of RNase activity, incorporation of RNA stabilizing modifications^57^, or alternative timing of readout, although such approaches fall beyond the scope of this study. Importantly, the absence of detectable eGFP signal does not imply that mRNA cannot be delivered into these cells. It may indicate that under the tested conditions, functional expression was not achieved at detectable levels.

Among the electroporation programs evaluated for BMDMs and HSCs, HSC-6 consistently yielded near-uniform RNP uptake and robust loss of target protein expression while preserving viability and differentiation capacity. We speculate that HSC-6 performed the best amongst the chosen electroporation programs because it delivers the highest amount of energy. Because mouse cells are smaller than their human counterparts, higher electroporation energy is required to open pores in the cell membrane. Typically, empirical testing of several electroporation conditions is required to find the most efficient electroporation program for a given cell type. With the MaxCyte ExPERT platform, the guesswork of testing dozens of combinations of voltage, pulse width, and other factors is taken out in favour of pre-programmed protocols tailored to a specific cell type. In this case, although no protocols for BMDMs existed, we predicted that because BMDMs are similar to HSCs, the electroporation protocols designed for human HSCs (HSC-4, HSC-6) would be effective for mouse HSCs and mouse BMDMs.

Electroporation can provide a non-viral delivery strategy that supports scalability and avoids unwanted genomic integration. The MaxCyte platform has been implemented in GMP-compliant clinical-scale electroporation using the ExPERT GTx flow electroporation system^58^, suggesting that the workflows described here can be readily adapted for large-scale translational applications. In addition, the workflows established here provide a modular framework in which rapid non-viral genome editing can be coupled with optional downstream viral delivery for reporter or rescue constructs, enabling flexible sequential gene disruption and re-expression while minimizing viral exposure.

Several limitations should be acknowledged. We inferred editing efficiency through protein loss and tracrRNA uptake rather than direct locus-level sequencing, and future studies incorporating DNA sequencing will enable more precise quantification of on-target efficiency and off-target outcomes. Additional validation using extended functional assays, including T cell effector activity, macrophage functional responses, and long-term HSC differentiation, will further strengthen translational relevance. Nonetheless, the strong preservation of viability, expansion capacity, and stress-response fidelity across lineages supports the robustness of the platform under the conditions examined in this study.

In summary, we establish MaxCyte ExPERT electroporation as a scalable, non-viral platform for efficient CRISPR RNP delivery across multiple primary mouse hematopoietic cells while preserving viability and lineage integrity. In activated OT-I T cells, both Exp T3 and Exp T4 programs supported robust delivery, with subtle activation-state–dependent tradeoffs in expansion at higher energy. In BMDMs and HSCs, we identify HSC-6 as a high-performing program for consistently high-level RNP delivery and target gene knockout. Together, these optimized workflows lower the barrier to rapid, reproducible genome engineering in primary hematopoietic cells for both discovery and translational applications.

## Acknowledgements

This work was supported by the following funding: a University of British Columbia 4-Year Fellowship (to J.H. Oh), BCCHR Canucks for Kids Fund Childhood Diabetes Laboratory Master’s Studentship (to D.Wi), BCCHR Jan Freidman Masters and Doctoral Studentships (to E. Tanaka), Breakthrough T1D—formerly JDRF—postdoctoral fellowship (3-PDF-2024-1504-A-N to L de Brito Monteiro), Michael Smith Health Research BC Trainee award (RT-2022-2619 to A-S Archambault), Banting postdoctoral fellowship (509775 to A-S Archambault), Canadian Institutes for Health Research (DT4-179512 to RI Klein Geltink), Natural Sciences and Engineering Research Council of Canada (RGPIN-2020-05390 to RI Klein Geltink), Canadian Cancer Society (Early Scholar Award to RI Klein Geltink), Michael Smith Health Research BC Scholar award (SCH-2022-2767 to RI Klein Geltink), University of British Columbia Department of Pathology and Laboratory Medicine and BC Children’s Hospital Research Institute startup funding (to RI Klein Geltink), and Canucks for Kids Diabetes Labs Legacy grant (to RI Klein Geltink).

## Author contributions

J.H.O., L.F.Y., E.T., and R.I.K.G. conceived the study. J.H.O., L.F.Y., E.T., D.W., L.M., A-S.A., and R.I.K.G. performed experiments and analyzed data. J.H.O., L.F.Y., E.T., L.M., D.W., and R.I.K.G. wrote the manuscript.; All authors reviewed the manuscript.

