## Supplementary Figures for "Versatile electroporation protocols enable reproducible CRISPR-RNP delivery across multiple primary mouse cells of the hematopoietic lineage"

Oh *et al.*

*Ramon I. Klein Geltink.

**This PDF file includes:**

Supplementary Figures 1 to 3

Supplementary Figure 1

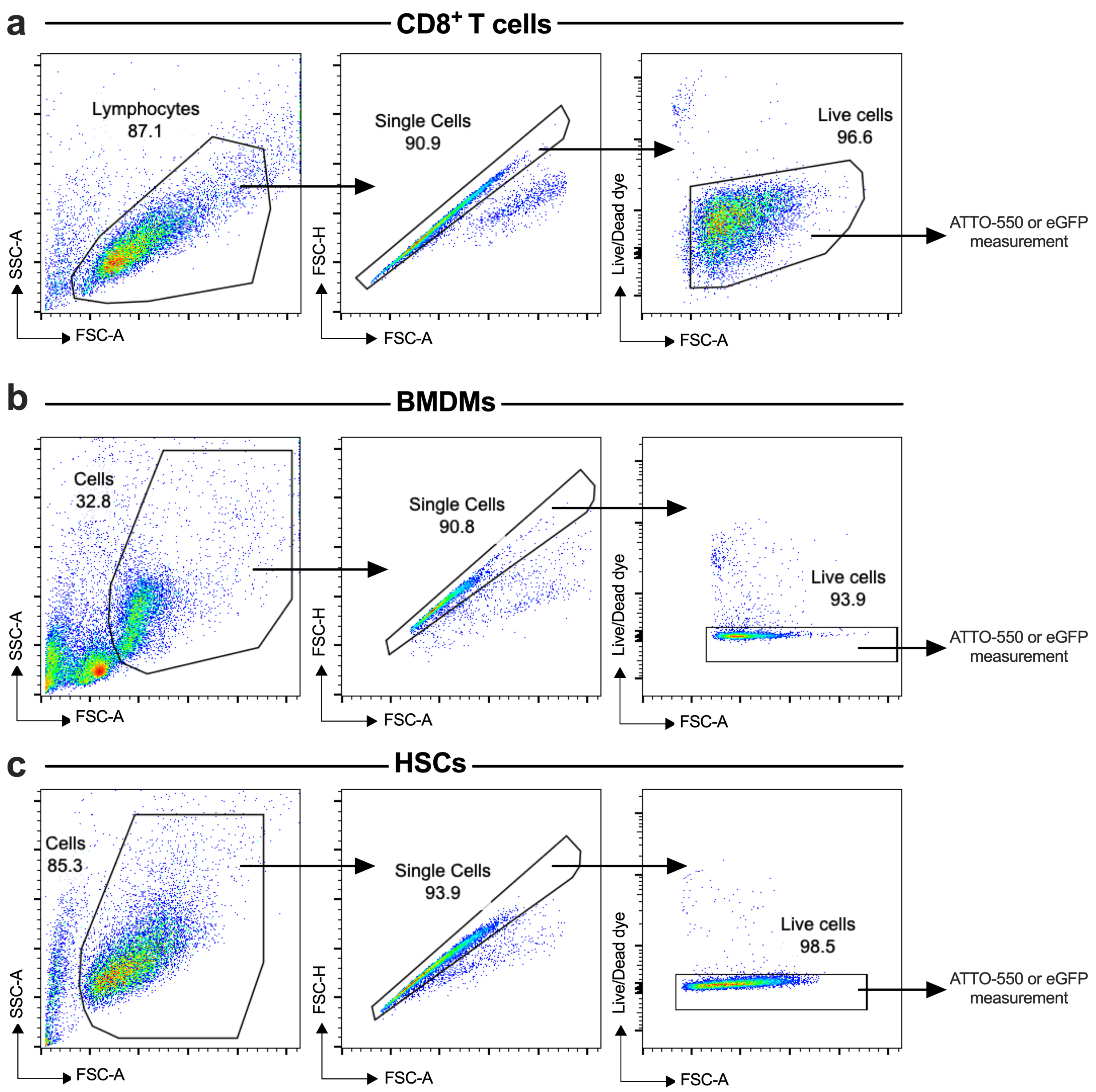


**Supplementary Fig. 1 Flow cytometry gating strategy for OT-I T cells (a), BMDMs (b), and HSCs (c).**

Supplementary Figure 2

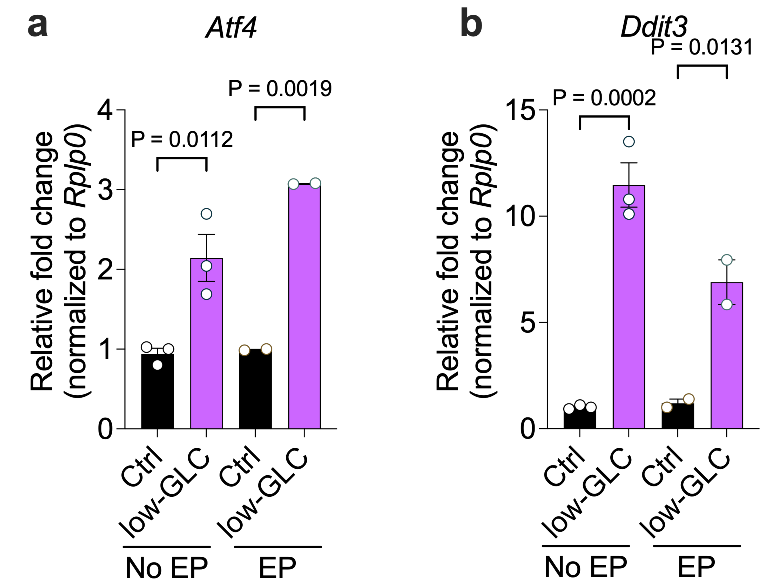


**Supplementary Fig. 2 Electroporation does not exacerbate ER stress marker expression in OT-I T cells. a, b** Control (Ctrl) and low-glucose (low-GLC) OT-I T cells cultured under no electroporation (No EP) or electroporation (EP) conditions were harvested, and total RNA was isolated for real-time PCR analysis of *Atf4* (**a**) and *Ddit3* (**b**) mRNA expression. *Rplp0* was used as the endogenous normalization control. Statistical significance was assessed using a two-tailed Student's t-test. Data are presented as mean ± SEM from n = 2–3 biological replicates.

### Supplementary Figure 3

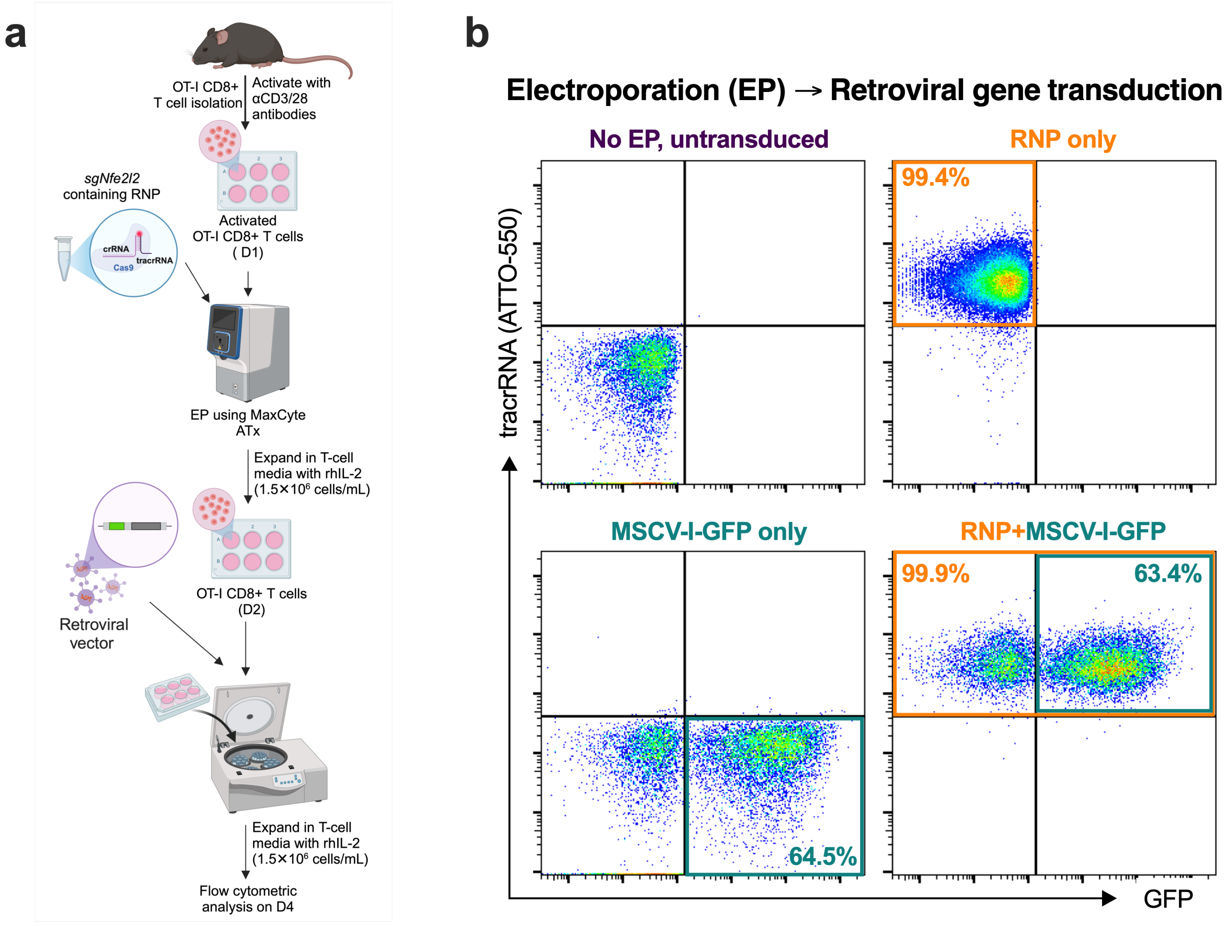


**Supplementary Fig. 3 Sequential electroporation and retroviral transduction enables efficient co-expression of CRISPR RNP and GFP reporters in OT-I T cells. a** Schematic depicting the experimental workflow combining CRISPR–Cas9 RNP electroporation and retroviral reporter transduction in OT-I T cells. On Day 1 (D1), OT-I T cells were activated for 24 hr with anti-CD3/CD28 antibodies in the presence of rhIL-2, followed by electroporation with RNP using the Exp T3 protocol. On Day 2 (D2), cells were retrovirally transduced with GFP reporter by spin-fection. Cells were subsequently expanded for an additional 48 hr in rhIL-2–containing medium and analyzed by flow cytometry. **b** Representative flow cytometry dot plots showing GFP fluorescence (x-axis) and ATTO-550–labeled tracrRNA fluorescence (y-axis), indicating successful retroviral reporter expression and RNP delivery following sequential electroporation and retroviral transduction.

Supplementary Figure 4

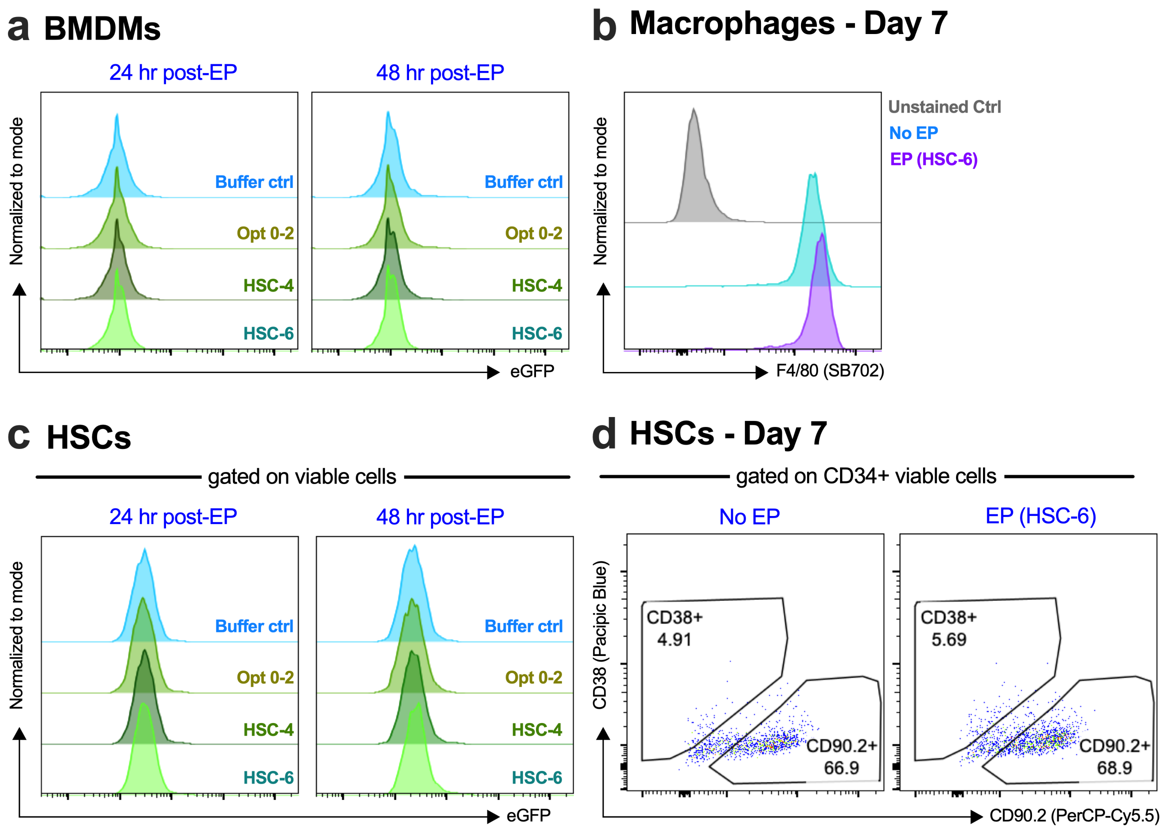


**Supplementary Fig. 4 Lack of detectable eGFP expression and preserved lineage marker expression in BMDMs and HSCs following electroporation. a** Representative histogram showing eGFP fluorescence in BMDMs at 24 hr (left) and 48 hr (right) following electroporation. **b** Representative histograms showing the frequency of differentiated macrophages ($F4/80^{+}$) in electroporated (EP) and non-electroporated (No EP) conditions. **c** Representative histograms showing eGFP fluorescence in HSCs at 24 hr (left) and 48 hr (right) following electroporation. d Representative flow cytometry dot plots showing the frequency of HSCs with a stemness phenotype ($CD38^{low}$ $CD90.2^{+}$) in EP and No EP conditions.
